# A Metagenomics – Based Diagnostic Approach for Central Nervous System Infections in Hospital Acute Care Setting

**DOI:** 10.1101/752618

**Authors:** Mohammad Rubayet Hasan, Sathyavathi Sundararaju, Patrick Tang, Kin-Ming Tsui, Andres Perez Lopez, Mohammad Janahi, Rusung Tan, Peter Tilley

**Author notes:** **Corresponding author:** Mohammad Rubayet Hasan, PhD, D(ABMM), FCCM, Assistant Professor of Pathology and Laboratory Medicine, Weill Cornell Medical College – Qatar, Clinical Molecular Microbiologist, Pathology Sciences Department of Pathology, Sidra Medicine, Level 2M, Office H2M-24093 PO BOX 26999, Doha, Qatar.

## Abstract

The etiology of central nervous system (CNS) infections such as meningitis and encephalitis remains unknown in a large proportion of cases partly because the diversity of pathogens that may cause CNS infections greatly outnumber available test methods. Here we present a metagenomic next generation sequencing (mNGS) based approach for broad-range detection of pathogens associated with CNS infections, which is suitable for application in the acute care hospital setting. Using an Illumina MiSeq benchtop sequencer and the IDseq pipeline for identifying pathogens in metagenomic sequence data, we show that the analytical sensitivity of mNGS to detect pathogens is comparable to that of PCR in simulated cerebrospinal fluid (CSF) specimens. We then applied this method for pathogen detection in 74 CSF specimens from patients with suspected CNS infections that were previously tested by culture and/or PCR. Diagnostic accuracy, sensitivity and specificity of mNGS approach with reference to conventional methods were all 95%. Furthermore, confirmatory testing on specimens that gave discrepant results were mostly in favor of the mNGS assay. The clinical application of mNGS holds promise to benefit patients with CNS infections of unknown etiology.

## Introduction

Central nervous system (CNS) infections such as meningitis and encephalitis are potentially life threatening diseases caused by a myriad of infectious pathogens. Besides high rates of mortality, meningitis and encephalitis are major causes of morbidity, and permanent disabilities such as brain damage, hearing loss, and learning disabilities can result from CNS infections (1–5). A specific etiologic agent cannot be identified in 15-60% of cases of meningitis and up to 70% of encephalitis (6, 7). Clinical management of meningitis and encephalitis is highly dependent on early and rapid detection of underlying causes of the disease, so that appropriate antimicrobial or anti-viral therapy can be instituted in a timely manner. Specific diagnosis is also important to avoid unnecessary treatment and hospitalization of patients with self-limiting forms of viral meningitis to minimize potential harm and unnecessary cost to patients (6).

Current diagnostic test methods for CNS infections include CSF Gram staining, CSF cell count, glucose, and protein measurements and biomarkers such as procalcitonin (PCT) and lactate. These tests are generally performed to distinguish between bacterial versus viral infections, and they are not specific for any causative pathogens. Bacteriological culture or PCR testing to detect specific pathogens in cerebrospinal fluid (CSF) are currently the most important methods for the diagnosis of CNS infections. However, a large number pathogens known to cause meningitis and encephalitis cannot be routinely cultured, and most molecular tests are targeted to common pathogens only. A broad-range, unbiased method to identify all pathogens in CSF would markedly improve the management of patients who are critically ill with undiagnosed disease.

Advances in genomic approaches—particularly in sequencing technologies—are being applied in many research and clinical settings. For example, next generation sequencing (NGS) technology, which is capable of deciphering millions of DNA and RNA sequences in parallel, has shown promise for detecting pathogens in clinical samples(8). In a recent online survey, with infectious diseases physicians, microbiologists and other associated professionals as participants, all respondents predict that NGS will find some use in the clinical microbiology laboratories within the next 5-10 years (9). Application of NGS in clinical microbiology laboratories include whole genome sequencing (WGS) of purified bacterial isolates for identification, typing, detection of antibiotic resistance, virulence profiling and epidemiological surveillance for infection control (9). On the other hand, NGS based metagenomic sequencing, which involves sequencing of all DNA content in the sample, has been applied mostly in research settings for microbiome studies as well as pathogen detection or characterization directly from clinical specimens (10–17). However, application of this technology in acute care diagnostic microbiology is limited due to its higher cost compared to conventional microbiological methods, lack of standardized methods, data interpretation challenges and the burden of analyzing and storing large datasets of sequences.

Recently, a heightened interest in the application of NGS in clinical microbiology laboratories has led to an increased number of studies aimed to optimize, standardize and validate mNGS for the diagnosis of CNS infections (18–23). The results of some of these studies are highly encouraging, particularly because of the ability of NGS to detect pathogens that are unidentifiable by conventional testing and their potential clinical impact in directing appropriate antimicrobial therapy. However, these methods are difficult to implement in acute care hospital settings because of complex laboratory workflows, too few specimens for batching, and lack of bioinformatics expertise for data analysis. We hypothesized that a simplified, low throughput approach adapted for implementation in an acute care diagnostic microbiology laboratory will provide actionable, clinically useful data with faster turnaround time and thus benefit a larger patient population with CNS infections of unknown etiology.

To this end, Illumina MiSeq is a low cost, benchtop sequencing platform that is suitable for applications such as small genome sequencing, target gene sequencing and 16S rRNA sequencing. MiSeqDx version of the instrument is also the first FDA-regulated, CE-IVD-marked, NGS platform for in vitro diagnostic (IVD) testing. MiSeq system offers simpler NGS library preparation protocol, very low input DNA requirement, high quality data and faster turnaround time. This platform is comparable to the larger Illumina HiSeq 2000 platform in its ability to sequence microbes from host-associated and environmental specimens(24). However, use of the MiSeq platform has not been standardized for metagenomic pathogen detection in clinical samples. In this study, we have determined that the analytical sensitivity of a MiSeq-based, shotgun metagenomic sequencing approach to detect bacterial and viral pathogens in cerebrospinal fluid (CSF) is comparable to PCR assays and then optimized a bioinformatic approach for high specificity based on a publicly accessible software platform for metagenomic detection of pathogens (https://idseq.net)(25, 26). The optimized approach was then applied to a set of retrospectively collected and previously tested CSF specimens (n=74) with an aim to describe the diagnostic accuracy, sensitivity and specificity of mNGS approach with reference to the standard methods. Our results suggest that mNGS-based testing can be implemented in clinical microbiology laboratories for the routine diagnosis of CNS infections, when broad-range detection of pathogens is required.

## Results

### Establishment of mNGS laboratory workflow

With an aim to establish a Illumina MiSeq based workflow for the detection of CNS pathogens, and to assess the analytical sensitivity of our mNGS approach, a set of CSF specimens (Training set) that were negative for CNS pathogens by routine laboratory investigations were spiked with a range of reference bacterial or viral strains at varying concentrations and assessed by both qPCR and mNGS. Negative CSF specimens (not spiked) and nuclease free water (NFW) were also assessed by mNGS to determine the level of background noise and contamination. Spiking was done keeping in mind that: i) the approximate titers of pathogens range from very low to medium to very high concentrations; ii) spiked organisms are typical CNS pathogens; and iii) spiked organisms represent various groups of bacteria and viruses such as gram positive and gram negative bacteria and enveloped and non-enveloped viruses. In order to reduce the cost of sequencing, multiple pathogens were spiked into the same specimen (Table 1). DNA concentrations in the CSF specimens, as well as the resulting NGS libraries prepared using the Nextera XT kit, were highly variable, but sufficient for sequencing (Supplementary Table 1). Also, the nucleotide (NT) reads associated with the internal control plasmid spiked into the specimens prior to extraction was highly variable with an average of about 3,070 reads per sample, ranging from 55 to 21,640 reads. However, the NGS reads aligned to pathogen genomes (NT reads) in IDseq were highly comparable to qPCR C_T_ values for the respective pathogens. While a statistically significant correlation (R= 0.63; p<0.001) was observed between approximate titers of spiked pathogens with their respective NT reads, the qPCR C_T_ values for different pathogens were more precisely correlated to their mNGS NT reads (R = 0.96; p<0.0001) (Figures 1 A and 1B). mNGS assay detected all spiked pathogens that were detectable by qPCR. Even a strain of *H. influenzae* that was spiked at a very low quantity, and thus undetectable by qPCR, was detected by mNGS. Based on these results, a simplified laboratory workflow was set for clinical validation (Figure 2).

**Table 1:** Composition, approximate titer, qPCR C_T_ values and mNGS sequence read outputs of spiked organisms in the simulated CSF specimens.

| Specimen no. | Organisms spiked | Approximate titer | qPCR result (C<sub>T</sub>) | IDseq NT reads |
| --- | --- | --- | --- | --- |
| Spike 1 | Human adenovirus type 7 | 4.7 TCID<sub>50</sub>/ml | 21.0 | 198,994 |
|  | Herpes simplex virus 2 | 3.7 TCID<sub>50</sub>/ml | 24.6 | 10,842 |
|  | *Haemophilus influenzae* | 5.9 CFU/ml | 27.7 | 83,901 |
|  | *Escherichia coli* | 5.9 CFU/ml | 29.2 | 21,971 |
|  | *Streptococcus pneumoniae* | 5.9 CFU/ml | 31.9 | 194 |
| Spike 2 | Herpes simplex virus 2 | 3.5 TCID<sub>50</sub>/ml | 22.3 | 121,258 |
|  | *Neisseria meningitidis* | 6.6 CFU/ml | 23.7 | 800,771 |
|  | *Streptococcus agalactiae* | 6.0 CFU/ml | 30.2 | 20,110 |
|  | *Escherichia coli* | 4.0 CFU/ml | 34.6 | 2,643 |
|  | *Haemophilus influenzae* | 2.0 CFU/ml | Undetermined | 48 |
| Spike 3 | Herpes simplex virus 2 | 3.5 TCID<sub>50</sub>/ml | 23.3 | 188,394 |
|  | *Neisseria meningitidis* | 6.6 CFU/ml | 24.2 | 1,744,667 |
|  | *Streptococcus agalactiae* | 6.0 CFU/ml | 32.6 | 12,761 |
|  | *Escherichia coli* | 4.0 CFU/ml | 34.3 | 6,825 |
|  | *Haemophilus influenzae* | 2.0 CFU/ml | Undetermined | 147 |
| Spike 4 | Herpes simplex virus 2 | 3.5 TCID<sub>50</sub>/ml | 23.2 | 141,614 |
|  | *Neisseria meningitidis* | 6.6 CFU/ml | 24.3 | 1,040,079 |
|  | *Streptococcus agalactiae* | 6.0 CFU/ml | 31.3 | 19,255 |
|  | *Escherichia coli* | 4.0 CFU/ml | 35 | 3,276 |
|  | *Haemophilus influenzae* | 2.0 CFU/ml | Undetermined | 88 |
| Spike 5 | Herpes simplex virus 2 | 3.5 TCID<sub>50</sub>/ml | 24.3 | 217,550 |
|  | *Neisseria meningitidis* | 6.6 CFU/ml | 25.1 | 1,720,187 |
|  | *Streptococcus agalactiae* | 6.0 CFU/ml | 33.9 | 5,787 |
|  | *Escherichia coli* | 4.0 CFU/ml | 35.4 | 6,825 |
|  | *Haemophilus influenzae* | 2.0 CFU/ml | Undetermined | 130 |
| NEG1 | None | 0 | - | - |
| NEG2 | None | 0 | - | - |
| NEG3 | None | 0 | - | - |
| NEG4 | None | 0 | - | - |
| NFW | None | 0 | - | - |

\*NFW, nuclease free water; NT, nucleotide; TCID, tissue culture infectious dose; CFU, colony forming unit

**Table 2:**
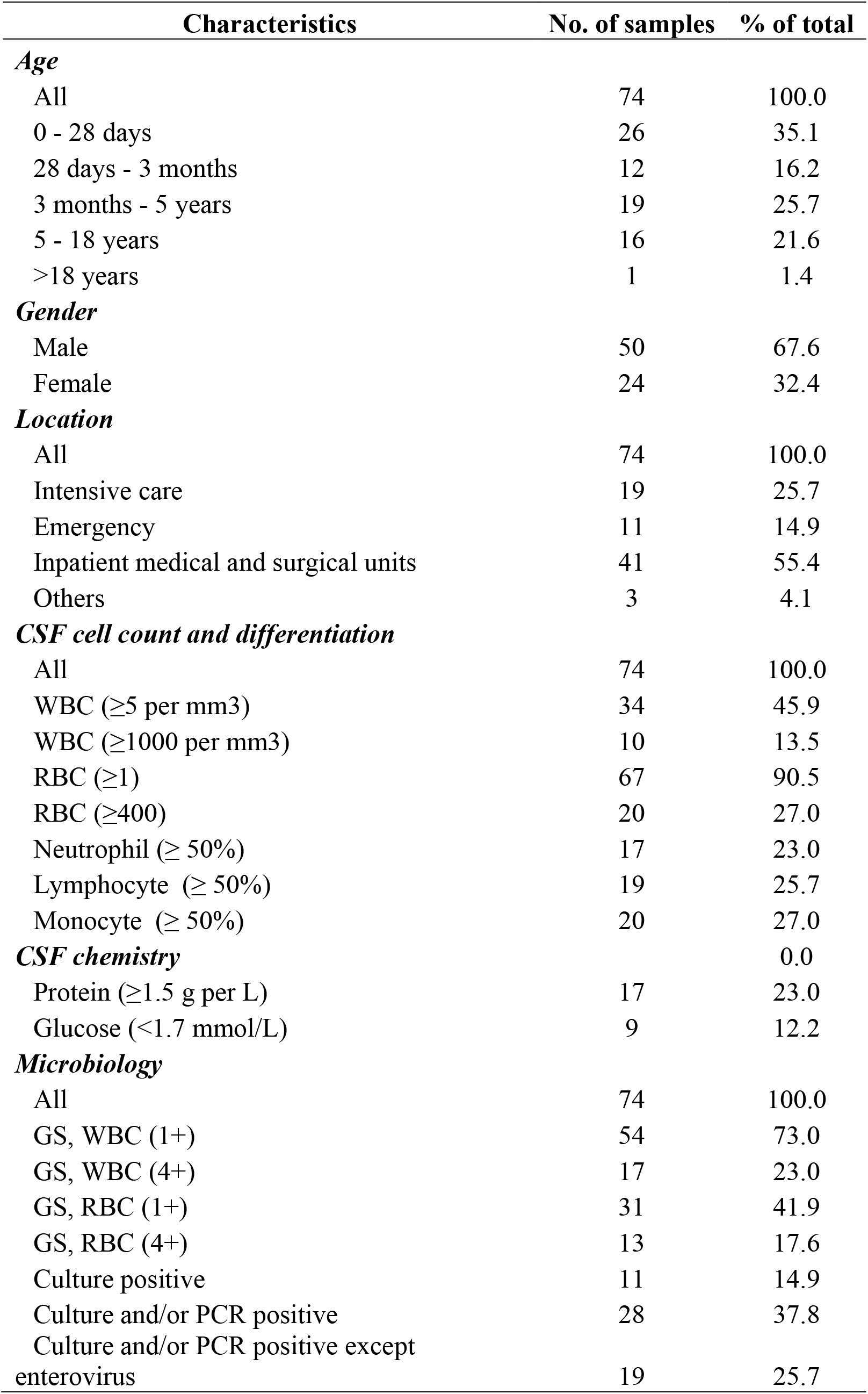
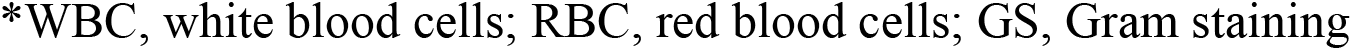
Specimen and patient characteristics.

| Characteristics | No. of samples | % of total |
| --- | --- | --- |
| ***Age*** |  |  |
| All | 74 | 100.0 |
| 0 - 28 days | 26 | 35.1 |
| 28 days - 3 months | 12 | 16.2 |
| 3 months - 5 years | 19 | 25.7 |
| 5 - 18 years | 16 | 21.6 |
| >18 years | 1 | 1.4 |
| ***Gender*** |  |  |
| Male | 50 | 67.6 |
| Female | 24 | 32.4 |
| ***Location*** |  |  |
| All | 74 | 100.0 |
| Intensive care | 19 | 25.7 |
| Emergency | 11 | 14.9 |
| Inpatient medical and surgical units | 41 | 55.4 |
| Others | 3 | 4.1 |
| ***CSF cell count and differentiation*** |  |  |
| All | 74 | 100.0 |
| WBC ($\geq 5$ per mm<sup>3</sup>) | 34 | 45.9 |
| WBC ($\geq 1000$ per mm<sup>3</sup>) | 10 | 13.5 |
| RBC ($\geq 1$) | 67 | 90.5 |
| RBC ($\geq 400$) | 20 | 27.0 |
| Neutrophil ($\geq 50\%$) | 17 | 23.0 |
| Lymphocyte ($\geq 50\%$) | 19 | 25.7 |
| Monocyte ($\geq 50\%$) | 20 | 27.0 |
| ***CSF chemistry*** |  | 0.0 |
| Protein ($\geq 1.5$ g per L) | 17 | 23.0 |
| Glucose ($< 1.7$ mmol/L) | 9 | 12.2 |
| ***Microbiology*** |  |  |
| All | 74 | 100.0 |
| GS, WBC (1+) | 54 | 73.0 |
| GS, WBC (4+) | 17 | 23.0 |
| GS, RBC (1+) | 31 | 41.9 |
| GS, RBC (4+) | 13 | 17.6 |
| Culture positive | 11 | 14.9 |
| Culture and/or PCR positive | 28 | 37.8 |
| Culture and/or PCR positive except enterovirus | 19 | 25.7 |

\*WBC, white blood cells; RBC, red blood cells; GS, Gram staining

**Figure 1:**
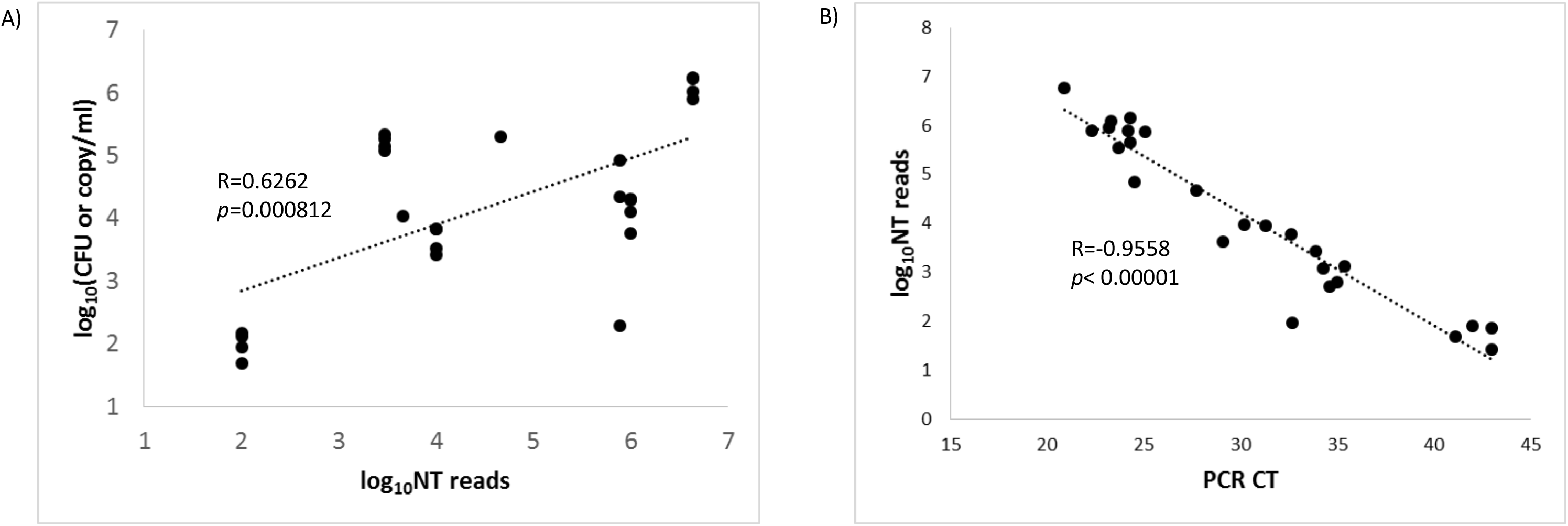
Analytical sensitivity of mNGS to detect pathogens in CSF specimens is comparable to qPCR. CSF specimens (n=9) negative by standard microbiological methods were spiked with a range of viral and bacterial pathogens at varying concentrations as described in the Materials and Methods or left unspiked and simultaneously tested along with a nuclease free water (NFW) sample by pathogen specific qPCR and by mNGS as described in the Materials and Methods. The approximate titer of pathogens (A) or qPCR C_T_ (B) values were plotted against mNGS read counts of pathogens obtained after analysis in IDseq.

**Figure 2:**
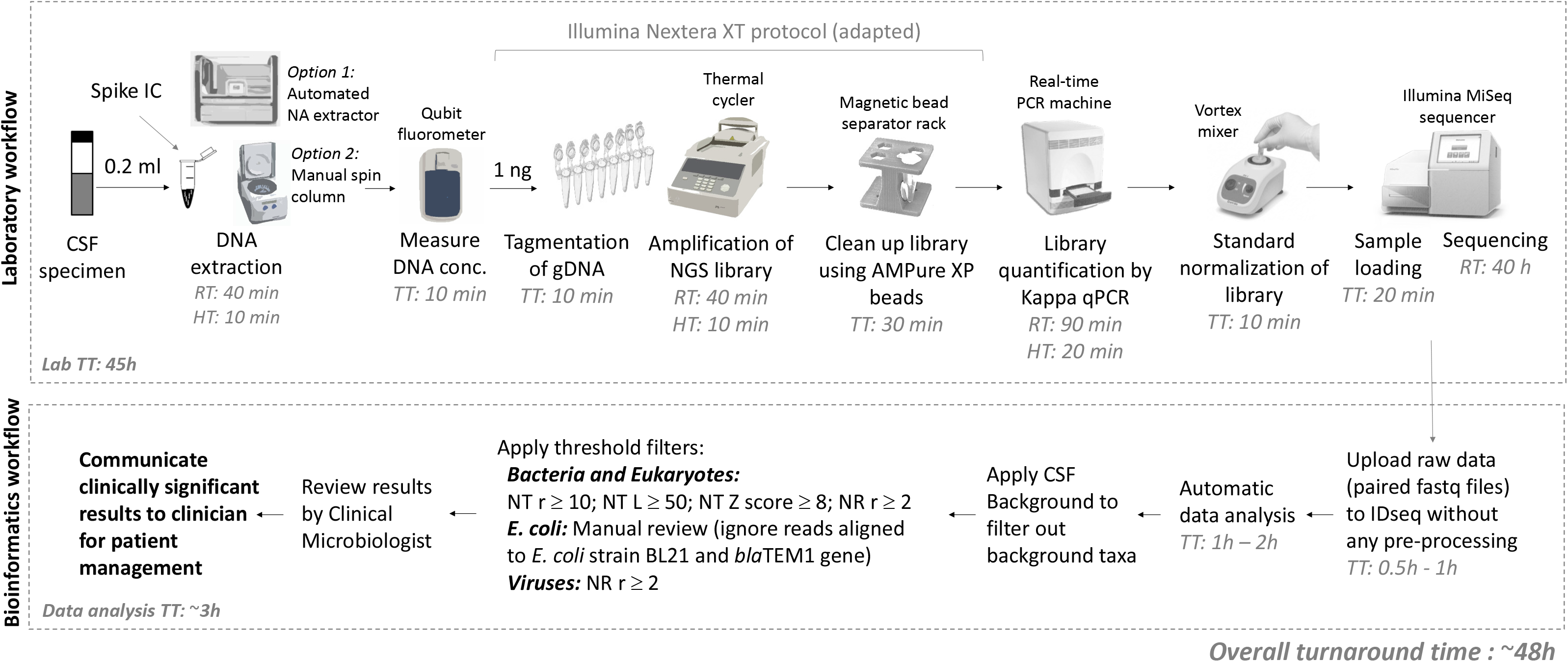
Schematic of mNGS laboratory and bioinformatics workflow. Total nucleic acids (NA) from CSF specimens were extracted after spiking pUC19 plasmid as an internal control (IC) on an automated extraction platform, Qiasymphony (Qiagen). However, any other extraction platform or manual, spin column based methods can also be used for DNA extraction. DNA concertation was determined using a Qubit fluorometer (Thermofisher). NGS library preparation and clean up were performed according to manufacturer’s instructions with the exception that 96-well plates were replaced by single tubes or PCR tube strips because single sample was processed. NGS library quantification, normalization, sequencing and data analysis were performed as described in the materials and methods. TT: total time, HT: hands-on-time, RT: reaction time, TAT: turnaround time. NT r : nucleotide reads, NT L : nucleotide alignment length in bp, NT Z score: nucleotide Z score, NR r : non-redundant reads.

### Establishment of mNGS bioinformatics workflow

While the DNA associated with all spiked pathogens were detected by mNGS using the IDseq software with high sensitivity, the specificity of mNGS results was poor using the default settings. In this setting, sequence reads from specimens mapped to a large number of taxa that include both common contaminants as well as bioinformatic artifacts originating from poor quality alignments making it difficult to differentiate true positive results from false positive results (Supplementary Figure 1). Along with the spiked specimens, this is also reflected in the IDseq results of known negative CSF samples as well as in the negative control water sample. Compared to the IDseq results, Metaphlan2 results were relatively more specific but had poor sensitivity. For example, Metaphlan2 failed to detect low concentrations of *S. pneumoniae* DNA and *H. influenzae* DNA in all samples. Even adenoviral DNA was undetectable despite being strongly positive by qPCR (Supplementary Figure 2). In order to improve the specificity of IDseq results instead, we first applied a background subtraction method. A background dataset was created from 40 negative CSF samples from the clinical validation study. The background dataset was then applied to the training set (n=10) and various filters were applied to adjust the sensitivity and specificity of IDseq results based on previous knowledge of spiked organisms and qPCR results. The optimal filter set that cleared all false positive results from the water sample and negative CSF samples were: i) NT reads ≥ 10; ii) NT Z score ≥ 8; iii) aligned base pair length ≥ 50 bp; and iv) Non redundant (NR) reads ≥ 2. While this approach significantly improved the specificity of mNGS (Supplementary Figure 3), true positive results for *E. coli* were filtered out because *E. coli* was also a common contaminant in mNGS. Therefore, we established a manual review process for *E. coli* results. We noted that contaminating *E. coli* DNA in our specimens mapped to *E. coli* strain BL21 or *E. coli* blaTEM1 gene and therefore these results can be ignored. *E. coli* results were only considered as positive if the NGS reads maps to a different strain of *E. coli* as the top hit. A phylogenetic tree of *E. coli* reads from the training set shows that *E. coli* DNA in most negative samples are closely related to either *E. coli* strain BL21 or *E. coli* blaTEM1 gene and all spiked specimens containing true *E. coli* DNA are closely related to NCBI reference genome *E. coli* CFT073. A review of individual sequence alignments in IDseq revealed that the only negative sample (Neg1) that clustered with *E. coli* CFT073 genome also has *E. coli* strain BL21 as its top hit (Supplementary Figure 4) and the number of sequence reads in this specimen aligned to *E. coli* CFT073 genome was much lower than BL21 (6 aligned reads versus 58 reads). We noted that unlike bacteria and eukaryotes, low count true viral reads were filtered out when the same filter set was applied. Therefore, to identify viral DNA in the test set data, we decided to apply a simple filter of only “NR reads ≥ 2” after background subtraction. All options for applying the background and customized filters are readily available in IDseq and are easy to select with few mouse clicks. The application of these customized filters in conjunction with the manual review for *E. coli* significantly improved the specificity of IDseq returned results without affecting sensitivity. However, the results should be interpreted by a clinical microbiologist to rule out: i) taxa that merely appear in the results because of sequence homology with the top hit; ii) clinically insignificant taxa; and iii) potential contaminant taxa introduced during specimen collection or handling and processing of specimens. A simplified bioinformatics workflow for clinical validation of mNGS is shown in Figure 2.

### Clinical validation of mNGS for pathogen detection in CSF

For clinical validation, previously saved residual CSF specimens (Test set; n = 74) that were submitted for microbiological assessment were selected for mNGS analysis. The samples were collected from patients with suspected CNS infections. The average age of patients was 3.3 years. 35% patients were neonates (≤28 days) and 68% were male. The neonatal and pediatric intensive care units and the emergency department accounted for about 26% and 15% specimens, respectively. 55% specimens came from other in-patient units including neuroscience and neurosurgery, respiratory and cardiac services and renal, endocrine and metabolic units. A review of patient’s laboratory data indicated that about 46% CSF specimens had abnormal white blood cell (WBC) counts and red blood cells (RBC) were detected in most of the specimens. The proportion of specimens with very high WBC (≥1000 per mm^3^) and RBC (≥400 per mm^3^) counts were 14% and 27%, respectively. Specimens considered to be “bloody taps” (RBC/WBC >500) were excluded. By Gram staining, about 23% and 18% specimens had very high (4+) WBC and RBC scores, respectively. By culture, 11 samples (15%) were positive and when combined with the PCR results 28 (39%) samples were positive for a bacterial, viral or fungal pathogen. A total of 9 samples were positive for enterovirus, which is an RNA virus. Because RNA viruses are not expected to be detected by the DNA-specific protocol described in this study, these samples were considered as negative samples for validation purposes.

The DNA concentration in CSF specimens ranged from 0.14 to 19.9 ng/μl with an average of 1.3 ng/μl (Supplementary Table 1). Because the Illumina Nextera XT protocol requires 1 ng DNA in 5 μl volume, 11 specimens had lower DNA concentration than the minimally required concentration. For these specimens, 5 μl of the undiluted DNA extracts were used for library preparation irrespective of their concentration. Other specimen extracts were diluted to 1 ng DNA as the starting material for library preparation. The NGS library concentration based on Kappa qPCR ranged from 0.4 to 150 nM with an average of 19.9 nM. The Illumina MiSeq protocol requires at least 2 nM DNA but 12 specimens had lower than the minimum library concentration. For these specimens, 5 μl of the undiluted libraries were processed for sequencing, while other libraries were diluted as required. A total of about 5 hours, with approximately 2 hours of hands on time, was required for DNA extraction, NGS library preparation, quantification, normalization and sample loading. Sequencing on the MiSeq required about 40 hours per run (Figure 2). The total sequence read output ranged from 3,757,836 – 44,424,176 with an average of 23,463,311. The average sequence read outputs obtained from specimens with <0.2 ng/μl DNA or with libraries with < 2 nM concentration (25,201,237 and 20,188,913, respectively) were not significantly different from the overall average sequence read outputs. Non-host reads ranged from 574 – 2,430,956 reads with an average of 56,179 reads, accounting for an average of 0.3% of total reads. The average run time for data analysis was 1.22 h. Data analysis for 90% and 74% of specimens were completed in <2 and <1 h, respectively.

By mNGS, performed according to the standard operating procedures described in Figure 2, a total of 19 samples were deemed positive for the presence of a pathogen (Figure 3). Compared to the conventional tests, mNGS missed one positive result (CW005), which was *Staphylococcus epidermidis* that had grown in enrichment broth only, and was originally reported as a potential contaminant. On the other hand, mNGS identified 4 additional pathogens that were not detected by conventional methods. *S. agalactae* was detected in two specimens from patients aged 0.09 and 0.03 years, respectively. The organisms were not recovered by culture, but Gram positive cocci were noted on Gram stain and were positive by confirmatory qPCR subsequently during the course of this study. In specimen CW060, mNGS detected few sequence reads that aligned to an *Acinetobacter* sp. TGL-Y2 plasmid, which was deemed clinically insignificant. A few reads of a *Streptococcus parasanguinis* were also detected in this specimen. A PCR targeting the bacterial 16S rRNA gene returned negative results for both specimens CW005 and CW060 (data not shown). Finally, low level HSV2 DNA was detected in sample CW322, which was strongly positive for *N. meningitidis*. Interestingly, our *E. coli* approach established based on training dataset correctly differentiated true positive results from false positive, background *E. coli* DNA, when applied to the validation dataset. A phylogenetic tree built based on the validation dataset clearly separated out *E. coli* DNA in 2 specimens that were related to the *E. coli* CFT073 genome from those that are related to the BL21 strain, and were interpreted as positive results (Supplementary Figure 5). It was noted that *E. coli* DNA in 69 out of 74 specimens were related to BL21 strains. The remaining 3 specimens were related to *E. coli* reference strain C9. However, all of these specimens were strongly positive for *N. meningitidis* and were therefore ignored. Among the viral taxa identified, Chimpanzee anellovirus and Torque Teno mini virus were deemed insignificant by the clinical microbiologist.

**Figure 3:**
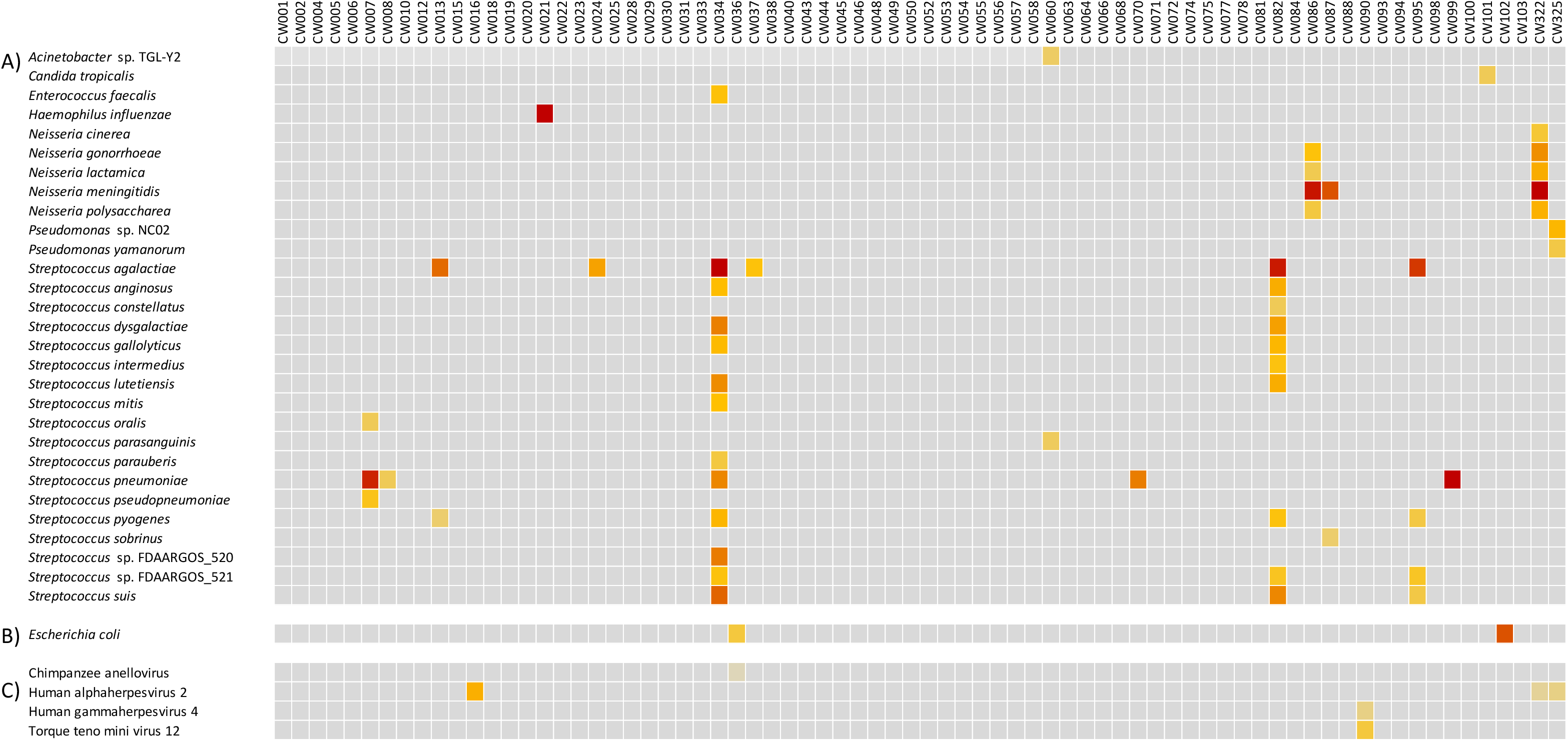
Heatmap of taxa identified in the validation set of specimens with IDseq after applying CSF background and customized threshold filters. mNGS was performed and data were analyzed as described in figure 2. Heatmaps were generated based on log_10_[NT total reads]. Different filter sets were applied for bacteria and eukaryotes versus viruses and *E. coli* reads were manually derived from IDseq after manual review of data.

Overall, the sensitivity, specificity and accuracy of mNGS results on 74 CSF specimens with the customized bioinformatic approach in IDseq were all 95% (Table 3). For comparison, all the mNGS data were also analyzed by the Metaphlan2 pipeline and the results were reviewed by a clinical microbiologist. All *E. coli* positive results, common environmental contaminants such as *Ralstonia picketii* and clinically non-relevant taxa were interpreted as negative results. The sensitivity, specificity and accuracy of mNGS results using Metaphlan2 were 58%, 96% and 86% respectively.

**Table 3:** Diagnostic performance of mNGS assay compared to conventional methods for CNS infections.

| Statistics | Metaphlan2 | IDseq |
| --- | --- | --- |
| Total number of results | 74 | 74 |
| True positive | 11 | 18 |
| True negative | 53 | 52 |
| False positive | 2 | 3 |
| False negative | 8 | 1 |
| Sensitivity | 58% (95% CI: 34%-80%) | 95% (95% CI: 74%-100%) |
| Specificity | 96% (95% CI: 87%-100%) | 95% (94% CI: 85%-99%) |
| Positive predictive value (PPV) | 85% (95% CI: 57%-100%) | 86% (95% CI: 67%-95%) |
| Negative predictive value (NPV) | 87% (95% CI: 80%-92%) | 98% (95% CI: 89%-100%) |
| Accuracy | 86% (95% CI: 77%-93%) | 95% (95% CI: 87%-99%) |

## Discussion

The potential for the application of NGS based, clinical metagenomics approach for the diagnosis of infectious diseases has been recognized for a few years now(8, 9, 15). Currently, initiatives are being taken to standardize the approach for routine application in the clinical microbiology laboratories(20, 22, 23). Because of the high cost and substantial expertise and labor involved, mNGS approach appears suitable only for specific applications in clinical microbiology such as for detection of pathogens in CSF for the diagnosis of meningitis and encephalitis because these infections are potentially life threatening and also because of the fact that a wide range of pathogens can cause such infections that are not routinely detected by standard methods. The present study was performed to establish and validate an mNGS approach for pathogen detection in CSF for the diagnosis of CNS infections that can be implemented in acute care hospital laboratories. The protocol has been developed for a low throughput setting so that single specimens can be processed without waiting for additional specimens for batching. Laboratory workflow and the data analysis workflow have been designed to be relatively simpler and faster compared to previously reported approaches. The procedure does not require huge capital investment or large scale instrumentation and automation, and can be performed by molecular microbiology technologists with a minimum of extra training. As with current culture methods however, the results must be reviewed by a clinical microbiologist for clinical correlation and to rule out false positive results arising from potential contaminants or clinically insignificant taxa.

Among the various platforms of Illumina that offers NGS at various depths and scales, the benchtop sequencer MiSeq was chosen because the equipment cost is relatively lower and the platform is more suitable for low throughput application. The MiSeq platform generates enough data and sequence reads so that low level pathogen DNA can be detected. Our analytical sensitivity data shows that the MiSeq-based approach was able to detect low level pathogen DNA with qPCR C_T_ 35 or higher. Even spiked *H. influenzae* that was undetectable by qPCR was detected by mNGS. Given the fact that PCR is widely considered to be one of the most sensitive method for pathogen detection, an equivalent or superior sensitivity to qPCR indicates that mNGS approach can be applied for the diagnosis of infectious diseases. In the clinical validation, we noted that despite minimal quality control and high variability in extracted DNA concentration, NGS library concentration, sequence read output, host versus non-host reads and IC reads, the mNGS approach was capable of identifying all pathogens but one that were identified by conventional methods. The only pathogen that was missed was a known skin contaminant that grew in enrichment broth only and was confirmed to be negative in this study. On the other hand, the fact that mNGS was able to identify *S. agalactiae* in two specimens that did not grow in culture suggests that mNGS approach is a powerful method for identification of fastidious pathogens or those that have failed to grow in culture because of prior antibiotic treatment.

Because NGS generates massive amounts of sequence data, interpretation of this data for clinical use is challenging. In order to develop a practical bioinformatics approach, with high sensitivity and specificity for pathogen detection in CSF, we tested various existing bioinformatics tools including Metaphlan2(27) and IDseq(28) using our training set. Metaphlan2 performs taxonomic assignment of metagenomic shotgun sequencing data at the species level using a unique database of clade specific marker genes as the reference database(27). Metaphlan2 is performed in command line, Linux or MacOS environments and therefore requires some bioinformatics expertise and experience working in a command line environment. IDseq is an online platform for metagenomic sequence analysis, where the user uploads raw sequence data to the platform and the rest of the procedure, including adapter trimming, data quality control, host DNA subtraction and alignment to both NCBI nucleotide (NT) database and non-redundant (NR) database is performed automatically.

When Metaphan2 was applied to our training and validation dataset, the specificity was high but the sensitivity was poor. For example, in the validation dataset, Metaphlan2 missed 8 of the 19 positive results by standard methods. On the other hand, when IDSeq pipeline was applied to our training data set, all spiked pathogens were detected with high sensitivity but the metagenomic sequence analysis returned a long list of bacterial, viral and eukaryotic taxa making it very difficult to differentiate true positive results from background taxa and/or potential contaminants, particularly when the target pathogen was present at low quantities (Supplementary Figure 1). The IDseq platform, however, offered us with a wide range of options to utilize additional statistical analyses and data filters to improve the sensitivity and specificity of the methods and allow for easier interpretation of data. First, we created a background dataset using data from 40 known negative samples. After background subtraction, while IDseq ranks taxa based on an aggregate score calculated from NT/NR ‘z scores’ and ‘reads per million’ (rpm), this method alone was not sufficient to delineate specific results from the list of potential pathogens and non-specific taxa. We therefore applied additional filters and manual review process as described in the results section in order to improve the interpretability of results. Interestingly, when this approach was applied to the validation data set, the returned results were highly accurate, sensitive and specific (95%) compared to the standard methods.

A limitation of our current approach is that we did not incorporate methods for detecting RNA viruses. A combined method for detection of both DNA and RNA targets by mNGS would expand the target pathogen range of the assay but would increase the cost of testing and increase the complexity of the procedure. Most pathogens detected by mNGS in our study were relatively common bacterial and viral pathogens, reflecting the nature of our pediatric population, but the high sensitivity and specificity of our approach suggests the potential for detecting more unusual pathogens in a higher risk population. We are currently undertaking a study to investigate the clinical impact of implementing mNGS for prospective pathogen detection in selected CSF specimens in a pediatric population.

In conclusion, we have developed a clinical metagenomic diagnostic approach for CNS infections which has demonstrated superior accuracy, sensitivity and specificity, compared to previously reported methods. The method offers faster turnaround time, minimal hands on time and can be easily implemented in an acute care diagnostic microbiology laboratory, with a moderate level of molecular expertise, and with modest capital investment. The cost of NGS is still much higher than standard microbiological methods. Therefore, application of this approach for routine CSF testing may not be economically viable at this time. Instead, the approach should be applied to selected patient specimens that are negative for a potential CNS pathogen by conventional testing, but where the clinical picture and CSF profile are highly suggestive of CNS infections. Currently, very few reference laboratories of the world offer clinical metagenomics based diagnostic services for infectious diseases. These laboratories are highly specialized and are equipped with large, production scale sequencing platforms and massive computational servers. The cost of sequencing is reduced through sample batching, but this, along with time required for sample shipping, increases the turnaround time, which can be detrimental for managing CNS infections. Therefore, the method described here, designed for use in a hospital acute care setting, should be of immediate benefit to patients with undiagnosed CNS infections.

## Materials and Methods

### Bacterial and viral strains

Bacterial strains used for spiking in this study were *Escherichia coli* (American Type Culture Collection [ATCC] 25922), *Streptococcus pneumoniae* (ATCC49619), *Streptococcus agalactiae* (ATCC12386), *Haemophilus influenzae* (ATCC10211) and *Neisseria meningitidis* (ATCC 13090) and viral strains include Herpes Simplex Virus 2 (HSV2) (ATCC VR-540) and Human Adenovirus type 7 (ATCC VR-7). *E. coli, S. pneumoniae* and *S. agalactiae* were grown on blood agar plates (Oxoid) overnight at 37°C in a 5% CO_2_ atmosphere. *H. influenzae* and *N. meningitidis* were grown on chocolate agar plates (Oxoid) under the same conditions. Viral stocks were maintained in Dulbecco’s Modified Eagle’s Medium (DMEM) supplemented with 10% DMSO at −80°C.

### Specimens

For test optimization and spiking, 9 CSF specimens submitted to the Microbiology and Virology laboratory of BC Children’s Hospital for culture and PCR testing between August 2013 and July 2014 were used. All CSF specimens were negative by culture and PCR for all pathogens that were used for spiking in this study. Following standard testing, residual specimens were kept at 4°C until processed. For clinical validation, residual CSF specimens (n=74) saved at −80°C after standard testing were used. These specimens were collected from August 2015 to December 2017 from children admitted to BC Children’s Hospital with suspected CNS infections. To maintain patient anonymity, all patient identifiers were removed by hospital staff, unaware of the current study results. Patient data including age and gender, ordering location, collection date, and laboratory data including CSF cell count and chemistry and Gram staining, culture and PCR results were recorded against a study specific identifier in a spreadsheet. Ethics approval for the study was obtained from the Research Ethics Boards of University of British Columbia, BC, Canada and Sidra Medicine, Doha, Qatar.

### Spiking

Bacterial suspensions were freshly prepared in phosphate buffered saline (PBS) to a turbidity equivalent to a 0.5 McFarland standard and further diluted 10, 100 and 1000-fold in PBS, as required. *N. meningitidis* (4.3 × 10^8^ CFU/ml), HSV2 (2.8 × 10^5^ TCID_50_/ml) and adenovirus (2.8 × 10^6^ TCID_50_/ml) were directly added from cultured stocks. Bacterial and viral preparations were spiked into 0.5 ml of 5 CSF specimens to give approximate final titers shown in Table 1, and vortexed for 10 sec before extraction of DNA. Four CSF specimens were left unspiked to serve as negative control.

### qPCR and DNA sequencing

Residual clinical samples were extracted on a QIAsymphony instrument (Qiagen) using the DSP Virus/Pathogen Mini kit. Total nucleic acids from spiked or unspiked specimens were extracted and analyzed by qPCR assays for various pathogens as described previously(29) and DNA concentration was measured in a Qubit 2.0 fluorometer using the Qubit dsDNA HS assay kit (Thermo Fisher Scientific, Inc.). To serve as an internal control for extraction, qPCR and NGS, a purified plasmid DNA pUC19 was spiked to each specimen prior to extraction at a final concentration of 1.4 × 10^5^ copies/ml. NGS libraries were prepared from 1 ng of extracted DNA using Nextera®XT DNA Sample Preparation Kit (Illumina), and sequencing was performed on an Illumina MiSeq sequencer using MiSeq Reagent Kit v2 (500-cycles) (Illumina) as described previously.(29) The concentration of prepared NGS libraries were determined by Kappa qPCR according to manufacturer’s instructions (Roche). Sequencing was performed in 3 different facilities: a) McGill University and Génome Québec Innovation Centre, Montréal (Québec), Canada b) Sidra Medicine, Doha, Qatar and c) Alliance Global Middle East, Beirut, Lebanon. Specimens were sequenced irrespective of the quality of NGS libraries. Confirmatory qPCR for *S. agalactiae* were performed as described previously.(29) Bacterial 16S rRNA PCR was performed as described previously.(30)

### Bioinformatics

Raw sequence data were analyzed either by Metaphlan2 pipeline(27) or uploaded to https://idseq.net for automated metagenomic analysis without any pre-processing of data. IC reads were obtained by mapping raw sequence reads to pUC19 plasmid sequence using Bowtie2 plugin in Geneious 11.1.5 software.

### Statistical analysis

The linear correlation of NGS data with that of PCR results was determined by calculating Pearson product-moment correlation coefficient in excel followed by determining the significance level at p<0.05. Diagnostic sensitivity, specificity, positive predictive value (PPV), negative predictive value (NPV) and accuracy (concordance) were calculated using an online, diagnostic test evaluation calculator, and associated 95% confidence intervals (CI) were calculated by the Clopper-Pearson interval or exact method using the same calculator.(31)

## Acknowledgements

The study was supported by British Columbia Children’s Hospital Foundation, Canada (Grant no. KRZ28061 to M.R.H.) and Sidra Medicine, Qatar (Grant no. SIRF_200026 to M.R.H.). We are grateful to the IDseq team for providing technical support for data upload and analysis.

